## Supplementary figures and images for "Functional hierarchies in brain dynamics characterized by signal reversibility in ferret cortex"

### Supp. Figure 1

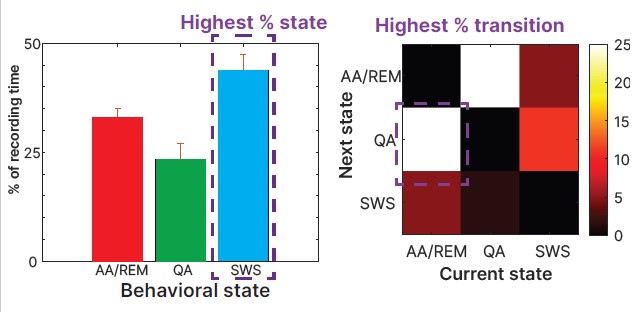
